# Genes in Humans and Mice: Insights from Deep learning of 777K Bulk Transcriptomes

**DOI:** 10.1101/2024.04.01.587517

**Authors:** Zheng Su, Mingyan Fang, Andrei Smolnikov, Fatemeh Vafaee, Marcel E. Dinger, Emily C. Oates

**Affiliations:** School of Biotechnology and Biomolecular Sciences, Faculty of Science, The University of New South Wales, Sydney, NSW 2052, Australia; BGI Research, Shenzhen 518083, China; BGI Australia, Herston, Queensland, Australia; School of Life and Environmental Sciences, Faculty of Science, University of Sydney, Sydney, NSW 2006, Australia

## Abstract

Mice are widely used as animal models in biomedical research, favored for their small size, ease of breeding, and anatomical and physiological similarities to humans^1,2^. However, discrepancies between mouse gene experimental results and the actual behavior of human genes are not uncommon, despite their shared DNA sequence similarity^3–8^. This suggests that DNA sequence similarity does not always reliably predict functional similarity. On the other hand, RNA-level gene expression could offer additional information about gene function^9,10^. In this study, we undertook characterization and inter-species comparison of human and mouse genes by applying innovative deep learning methodologies to a large dataset of 410K human and 366K mouse bulk RNA-seq samples. This was achieved by using gene representations from our Transformer-based GeneRAIN model^11,12^. These gene representations aggregate information from large gene expression datasets, and provide insights beyond DNA sequence similarity. We identified 2,407 human-mouse homologous genes with high DNA similarity but distinct RNA characteristics, and showed that these genes are more likely to have differing disease/phenotype associations between the two species. Additionally, we found 3,070 homologous genes with low similarity at both the DNA and RNA levels, suggesting the highest risk of discrepancies in study results between the two species. We propose that this approach will support future decision making around whether the mouse will be an appropriate model for studying specific human genes, and whether the results of specific mouse gene studies are likely to be recapitulated in humans. Our methodological innovations offer valuable lessons for future deep learning applications in cross-species omics data. The interspecies gene relationship findings from our study also contribute valuable insights into the gene biology and evolution of the two species.

## Introduction

Mouse models are extensively used in various human biomedical research scenarios, including the study of disease mechanisms, validation of gene functions, and drug development^1,2,4,6,13^. The sequencing of both human and mouse genomes has dramatically increased our knowledge of gene relationships between these two species. It has enabled us to identify mouse homologous genes that could represent human genes using DNA sequence similarity information^14–16^.

However, mouse genes selected based on DNA sequence similarity do not always mirror the function of human genes. Studies have shown discrepancies between outcomes in mouse models and human clinical settings^3–8^. These inconsistencies might arise from cellular, organ-level, or environmental differences between humans and mice. They might also stem from the limitations of using DNA sequences to infer gene function similarities. DNA provides a static blueprint of biology but may not fully capture the complexity of gene function, as genes typically function via the RNAs and proteins they encode^17^. RNAs may therefore offer an additional layer of information about gene function.

Although multiple mouse gene expression studies have been conducted, they have often relied on techniques such as dimension reduction, phylogenetic clustering, co-expression analysis, and differential expression analysis^16,18–33^. These traditional methods face challenges in achieving a comprehensive comparison of genes at the RNA level, as they can be susceptible to batch effects, biased by small sample sizes, and constrained by the limited availability of samples from matched biological conditions^33,34^. Given the dynamic and complex nature of gene expression, which varies across genders, ages, tissues, and conditions, a thorough characterization at the RNA level necessitates integrating data from diverse biological contexts and a large collection of samples.

In this study, our goal was to characterize the similarities and differences between human and mouse genes at the RNA level. This was achieved by integrating information from 410K human and 366K mouse bulk RNA-seq samples. Our approach leveraged advanced deep learning methods, including state-of-the-art Transformer models^11^, self-supervised techniques^35–37^, and the development of embedding alignment methodology.

## Results

### Representing genes across species using RNA information

In our study, we started by creating representations of protein-coding genes, by integrating gene expression information from 410,850 human and 366,750 mouse bulk RNA-seq samples from the ARCHS4 database^38^. These samples were from a wide range of tissues, genders, ages, and biological conditions. Our gene representations were created using the ARCHS4 RNA-seq datasets to train our state-of-the-art gene representation GeneRAIN model^12^, a Transformer-based^11^ Generative Pre-Training (GPT) model^37^. The gene representations were 200-number vector embeddings from the model. Prior study revealed that the gene embeddings generated by GeneRAIN model could effectively represent genes by encoding a comprehensive set of biological information^12^. This includes information about gene-associated diseases/phenotypes, protein interactions, transcription factors, biological pathways, gene ontology, associated cell types and more. Throughout this text, we will use the terms ‘gene embedding’ and ‘gene representation’ interchangeably.

In the study, we performed intensive methodological development and evaluation to represent genes of the two species in the same space. Here, a ‘space’ is a mathematical framework where similar numerical values in the gene representations represent similar biological attributes. Conversely, if genes are represented in separate spaces, the same numerical values can represent different attributes. Traditional single-species models will represent genes of two species in distinct spaces, making comparison challenging. Thus, for this study, developing methods to represent genes from two species in the same space was crucial to make them comparable.

We started with a mixed training approach, inspired by joint training techniques used in aligning cross-lingual word embeddings^39^. In this approach, human samples were randomly mixed with mouse samples to create a training dataset for a single GeneRAIN model (Fig. 1a). In the model, 18,757 human genes and 19,163 mouse genes had their own gene embeddings but shared other model parameters. Throughout the training, although the model progressively ‘pulled’ the gene embeddings of the two species closer (Extended Data Fig. 1a and 2, observe the percentage of explained variance on the PCA plot x-axis), they were not in a same space (Fig. 1d), indicating the need for further methodological enhancements.

**Fig. 1.**
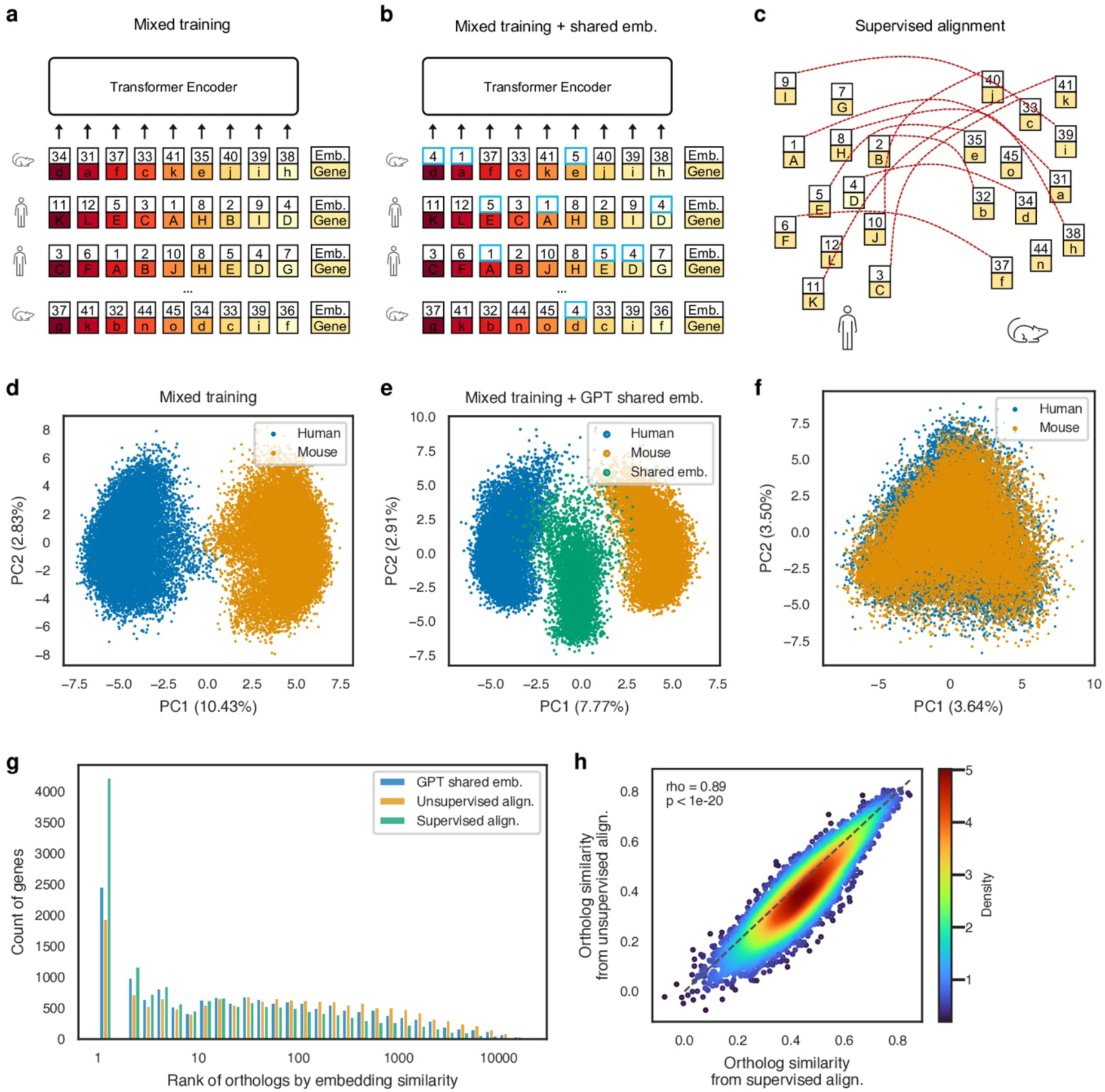
Alignment of cross-species gene embeddings. **a,** Schematic of mixed training: Human RNA-seq samples were randomly mixed with mouse samples before being fed into a single Transformer model. Four samples are illustrated as examples. Lower boxes in each line denote gene symbols, while upper boxes indicate embedding indices. The colors represent gene expression level. Genes were ordered from highest to lowest expression prior to model input. Upper- and lower-case gene symbols represent orthologs in the two species. Different embeddings were used for genes in two species. **b,** Schematic of mixed training plus shared embedding strategy. Some randomly selected one-to-one orthologs from both species were forced to share identical embeddings (denoted by blue boxes). **c,** Supervised alignment approach. One-to-one orthologous gene pairs (connected by dashed lines) were used to supervise the alignment of gene embeddings from two species. **d to f,** PCA plots depicting the distribution of gene embeddings from three different alignment strategies, with each dot symbolizing a gene. **g,** Distribution of embedding similarity ranks for orthologs. For each human gene, the embedding similarity rank of its ortholog (where available) among all mouse genes was determined, and this histogram displays the distributions of these ranks from three different embedding alignment strategies. **h,** Correlation of orthologous gene similarities from the supervised alignment and unsupervised alignment approaches. This scatter plot visualizes the Spearman correlation of cosine similarities of one-to-one orthologous gene embeddings, from supervised alignment and unsupervised alignment approaches. The dashed line represents the line of equality (y=x).

Subsequently, we adopted a ‘shared embeddings’ approach to further align gene embeddings, using the information of 16,983 one-to-one orthologous genes from the MGI database^40^. As a significant portion of these orthologous genes should share similar functions across the two species^41^, they should have very similar gene representations. Thus, we forced the 5,000 randomly selected one-to-one orthologous gene pairs to have the same gene embeddings (Fig. 1b). We hoped to use these 5,000 gene pairs as anchors to further pull the representations of other genes closer together. Note that only the representations of these other genes would be used for later analyses. Three different models, each applying shared embeddings to a different set of orthologs, were trained to make all genes have usable embeddings (Methods). Our analysis indicated that, although this method did bring the gene embeddings closer together, they remained in relatively distinct spaces (Fig. 1e).

Despite being in distinct spaces, we investigated whether the gene embeddings already encoded information about inter-species gene relationships. This was done by examining if orthologous genes from the two species had similar representations. It was found that, for 2,459 of the 16,983 human gene examined, their mouse orthologs were the nearest embedding neighbor among all mouse genes. Additionally, for 6,007 human genes, their mouse orthologs were among the ten nearest embedding neighbors (Fig. 1g). This suggests that the gene embeddings had indeed captured some inter-species gene relationship information.

We further explored representation alignment strategies by testing the advanced unsupervised and supervised embedding alignment methods from MUSE^42^ (Fig. 1c), on the embeddings from mixed training. These methods adopt approaches of contrastive learning and Procrustes alignment^43^, and find a single transformation matrix to align all the genes. Assessment revealed that unsupervised method had limited efficacy (1,932 orthologs being the nearest, Fig. 1g). However, the supervised method successfully made 4,216 human orthologs the nearest neighbor, and 8,140 human orthologs ranked among the top 10 closest embeddings (Fig. 1f, g).

Additionally, to evaluate how the result can be impacted by different alignment techniques, we assessed the correlation of embedding similarities from different approaches, focusing on one-to-one orthologous gene pairs. Strong correlation was observed between supervised alignment and unsupervised alignment approaches, especially in the high similarity orthologs (Spearman correlation rho = 0.89, p < 1e-20, Fig. 1h). The high agreement with the unsupervised approach indicates low risk of overfitting in the supervised approach. We also compared similarities between shared embeddings and supervised alignment approaches. Despite lower overall similarities from the shared embedding approach, strong correlation was observed (Spearman correlation rho = 0.74, p < 1e-20, Extended data Fig. 3), indicating gene relationship results were robust to the used embedding alignment techniques.

To further optimize and evaluate embedding alignment methodologies, we performed multiple experiments, as detailed in Supplementary Information. Briefly, it was revealed that combining shared embeddings with supervised alignment methods slightly further enhanced the performance. Our ‘positive control’ experiment demonstrated the high effectiveness of our embedding alignment approach in reconstructing inter-species gene relationships. The experiment indicated that it was not due to methodology limitations that fewer than 5,000 of 16,983 mouse genes were the nearest embedding neighbors of their human orthologs. Additionally, a ‘negative control’ experiment confirmed that our approach did not arbitrarily align genes in the supervision set regardless of their embeddings. We also showed the alignment results were not limited by training epochs or number of orthologous gene pairs for supervision, and they were robust to Transformer architectures. Moreover, mixed training proved beneficial for embedding alignment.

To mitigate bias from a single approach, we averaged the gene pair similarities from shared embeddings plus supervised alignment approach and supervised alignment-only approach. We used the averaged similarities for subsequent analyses. Our comprehensive optimization and evaluation of embedding alignment methodologies created a solid, reliable foundation for future analysis and application of these gene representations.

### RNA embeddings and DNA attributes in homologous genes

With human and mouse gene representations learned from expression data in a shared space, we analyzed inter-species gene similarities. We observed varying degrees of representation similarity (termed as RNA similarity or RNA simi.) among human-mouse homologous gene pairs. To facilitate comparison, for each pair, we determined the rank of the mouse gene based on its similarity to the corresponding human gene, compared to all mouse genes. We then categorized these homologs into two groups based on their ranks: a high similarity group comprising the top 40% of homologs (n = 7,249, rank <= 3) and a low similarity group including the bottom 40% (n = 7,103, rank >= 21). We excluded the middle 20% to avoid ambiguity in the similarity property.

We explored the DNA attributes of two homolog groups. We examined DNA sequence similarities, which we termed as DNA similarity or DNA simi., in coding regions, as well as 1kb upstream and downstream regions of transcription sites through pairwise transcript level alignment. Our analysis revealed that homologs with high RNA embedding similarity generally exhibited high DNA similarity in all these regions, though the overall DNA similarity was lower in the non-coding regions (Methods, Fig. 2a-c, Extended Data Fig. 6a-c and Supplementary Table 1-4). We also noted that homologs with high RNA similarity have higher GC content (Fig. 2d and Extended Data Fig. 7a-e), indicating stronger guanine-cytosine base pairs and greater molecule stability^44^. Furthermore, high RNA similarity homologs tended to have slightly longer coding sequences (Fig. 2e and Extended Data Fig. 6d).

**Fig. 2.**
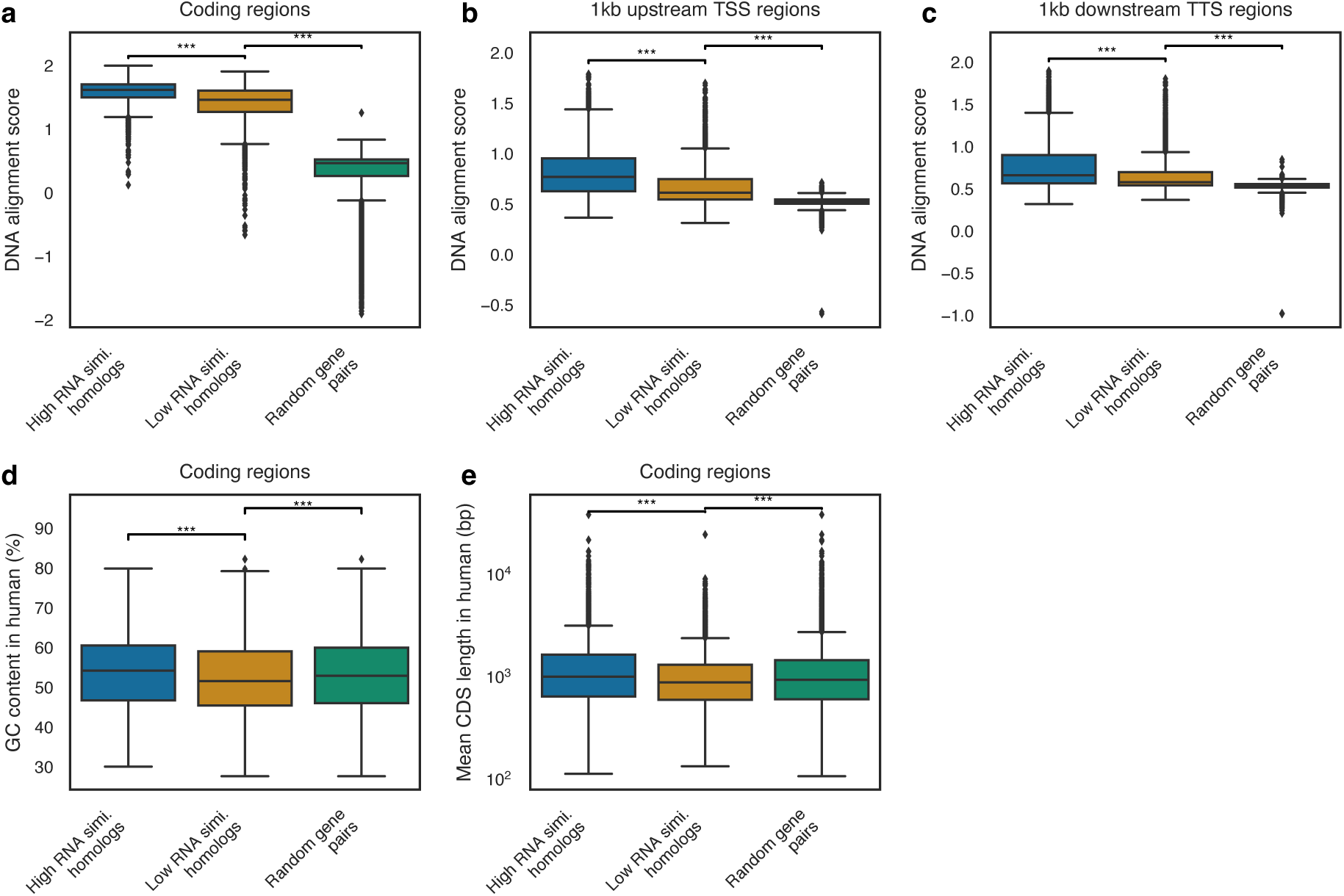
DNA characteristics of homologs grouped by embedding similarity. **a-c,** Boxplots depicting DNA alignment scores (Methods) for **(a)** coding sequences, **(b)** 1kb upstream regions of transcription start sites, and **(c)** 1kb downstream regions of transcription termination sites in homologs with high embedding similarity (n = 7,249) and low embedding similarity (n = 7,103). Random gene pairs (n = 29,998) are included for comparison. DNA alignment scores are determined by the highest pairwise alignment similarity across transcripts for each gene pair, reflecting the extent of DNA sequence conservation. **d,** GC content in the human genes from three gene pair groups. **f,** Average coding sequence length in the human genes from three gene pair groups. In all boxplots: center line, median; box limits, upper and lower quartiles; whiskers, 1.5× interquartile range; outliers, points. All comparisons were evaluated using a two-sided Wilcoxon rank-sum test. *** *p* < 0.00001, ** *p* < 0.001, * *p* < 0.05, ‘ns’ not significant (*p* >= 0.05).

### RNA similarity and phenotype associations

Discrepancies between human and mouse gene study results could arise because human and mouse homologs do not share the same disease or phenotype associations. We explored whether gene representations derived from RNA information could better explain phenotype association than DNA sequence similarity. For this purpose, homologs were further categorized into high (top 40%, based on coding region DNA alignment scores) and low (bottom 40%) DNA similarity groups. We identified homologs with high similarity in both DNA and RNA (n = 4,694), in only RNA (n = 1,107), in only DNA (n = 2,407), and in neither (n = 3,070).

For this investigation, we used mouse/human orthology phenotype annotations from the MGI database^40^ (Methods). Analysis revealed that homologs with high similarity in both RNA and DNA had the largest number of shared associated phenotypes. This was followed by homologs with high RNA but low DNA similarity, then those with low RNA but high DNA similarity, and finally, homologs with low similarity in both (Fig. 3a and Supplementary Table 2). This suggests RNA similarity provides information not always captured by DNA similarity. Moreover, homologs with low DNA similarity can have more shared associated phenotypes than those with high DNA similarity, if the former have higher RNA similarity. A similar pattern was observed when analyzing the percentage of homologs with shared associated phenotypes (Extended Data Fig. 8a).

**Fig. 3.**
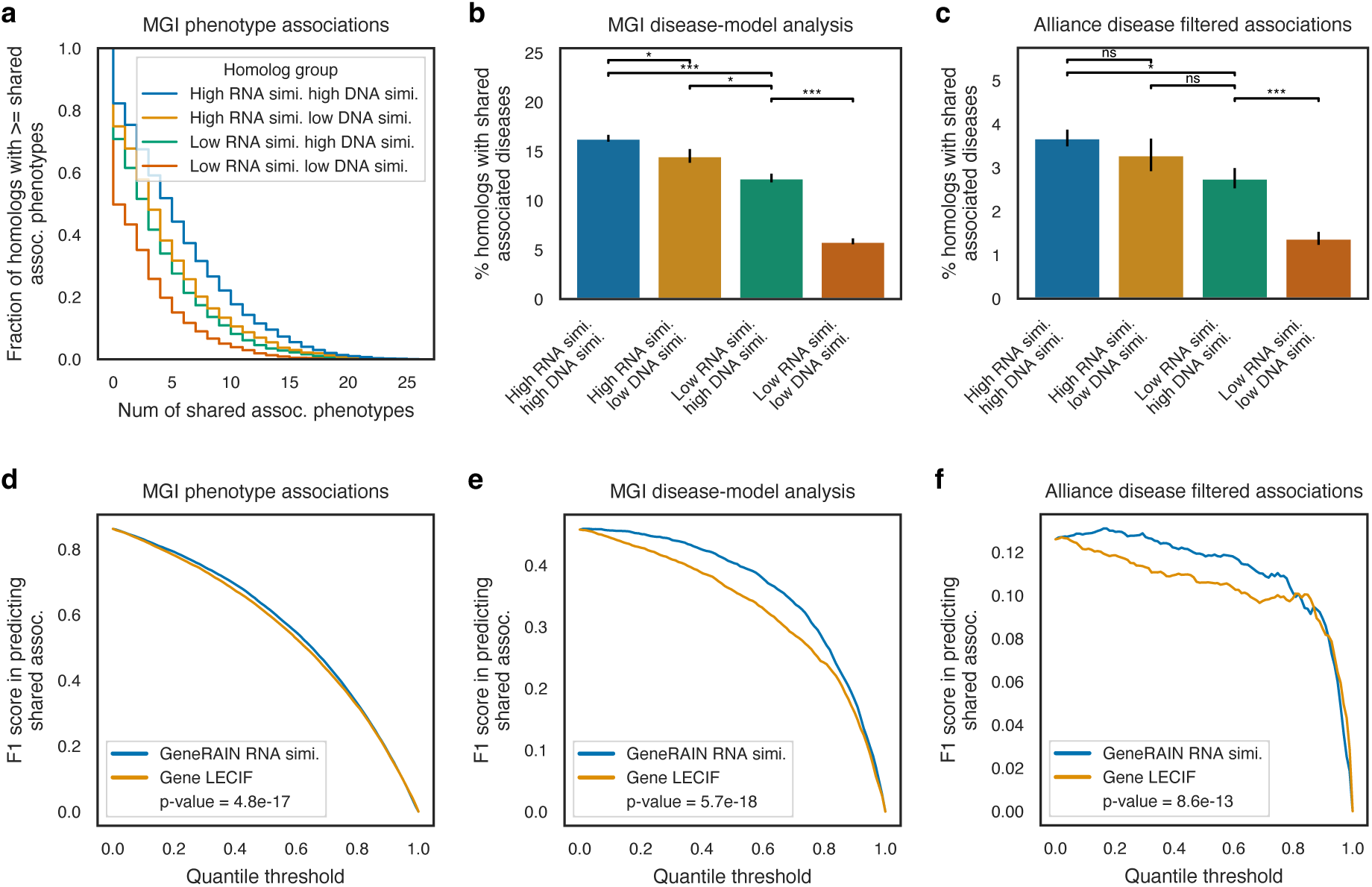
Embedding similarity and phenotype association analysis. **a,** Empirical cumulative distribution function (ECDF) plot illustrating the number of shared phenotype associations for homologs, categorized by DNA similarity and embedding similarity learned from RNA data. The horizontal axis represents the number of shared associated phenotypes for each homolog gene pairs, while the vertical axis denotes the fraction of homologs with a greater number of shared associated phenotypes. Lines higher on the plot indicate more shared associated phenotypes. All group comparisons had p-values less than 1e-14, except for the second vs. third group (p-value = 0.0039), two-sided Kolmogorov-Smirnov test. **b,** Barplot displaying the percentages of homologs with shared associated diseases, according to the human disease mouse model annotations from MGI database. **c,** Barplot showing the percentages of homologs with shared associated diseases, according to filtered disease association annotations from the Alliance of Genome Resources database (Methods). Error bars represent the standard errors in both barplots. In barplots, two-sample z-test for proportions was used for comparisons. *** p < 0.00001, ** p < 0.001, * p < 0.05, ‘ns’ denotes not significant (p ≥ 0.05). **d-f**, F1 Score vs. threshold curves comparing the performance of our RNA similarities and LECIF scores in predicting whether homologous gene pairs have shared association(s), using annotations from the database as indicated in the title. For LECIF, the average score within the genic region was used for each gene. The curves visualize how the F1 score (harmonic mean of precision and recall) varies across different classification thresholds. The p-values, obtained from the Wilcoxon signed-rank test, compare the F1 scores of the two metrics across 100 quantile thresholds.

To mitigate bias from using a single phenotype annotation dataset, we analyzed another dataset, specifically the ‘mouse models of human diseases’ annotations from the MGI database. This dataset includes information about human diseases, their mouse models, and the associated genes. Given the low number of shared associated diseases among the homologous genes (Extended Data Fig. 8b), we focused on comparing proportion of each gene group with shared disease association(s). Results indicated that homologs with high RNA similarity have largest proportion of shared association(s). In contrast, homologs with low RNA similarity showed a smaller proportion, even if their DNA similarity is high (Fig. 3b and Supplementary Table 2).

We extended our analysis using the third phenotype association annotations, specifically the Alliance of Genome Resources database^45^. As association type and evidence type information is available in this annotation dataset, we filtered the association annotations to retain only those with well-supported evidence (Methods). This analysis reaffirmed similar patterns of shared disease associations (Fig. 3c, Extended Data Fig. 8c and Supplementary Table 2), reinforcing the notion that RNA similarity offers additional, important information not captured by DNA sequence similarity.

We then compared the performance of our embedding similarities for phenotype association with an existing, state-of-the-art human-mouse gene conservation metric called LECIF^33^. LECIF assesses human-mouse genomic functional conservation by integrating epigenomic, transcription factor binding, and transcriptomic data using contrastive learning neural networks. It incorporates data on over 8,000 human and 3,000 mouse features from consortia like ENCODE^46^, Roadmap Epigenomics^47^, and FANTOM^48^. Our results showed that our representation similarities outperformed LECIF in predicting shared phenotype and disease associations of human-mouse homologs. (Fig. 3d-f, Extended Data Fig. 8d-f). This suggests that our approach, although relying solely on transcriptome data, can achieve superior performance compared to models that incorporate multi-omics data.

Given the predictive ability of our RNA embedding similarities, this approach could potentially help explain discrepancies observed in mouse model studies. For example, transgenic mouse models of certain genetic disorders may only partially replicate human disease phenotypes, or therapeutics targeting some genes or their encoding proteins may show promise in mice but fail to result in a positive clinical outcome in humans. Examples include discrepancies observed in studies of SOD1 (similarity rank 171; rank ≥ 21 was considered as low similarity), C9orf72 (rank 89) for Amyotrophic Lateral Sclerosis^49,50^, PSEN2 (rank 587) for Alzheimer’s Disease^51,52^, PARK7 (encodes DJ-1, rank 105) for Parkinson’s disease^8,53–55^, and IL4 (rank 528), NOS2 (rank 50), and PTAFR (rank 33) for asthma^8,56^. These genes have low RNA similarities in mice, suggesting that the correlation of these genes with other genes and possibly their functions have diverged between mice and humans. This divergence could contribute to the discrepancies observed between the human and mouse studies.

### Modelling lncRNAs and pseudogenes across species

Finally, we expanded our analysis to include 20,184 long noncoding RNA (lncRNA) genes and 8,439 pseudogenes alongside protein-coding genes in the model (Extended Data Fig. 1e). We employed mixed training and supervised alignment for the expanded geneset. We found that the similarities obtained were highly correlated with those from the protein-coding only model (Fig. 4a), indicating consistent performance when modelling more genes.

**Fig. 4.**
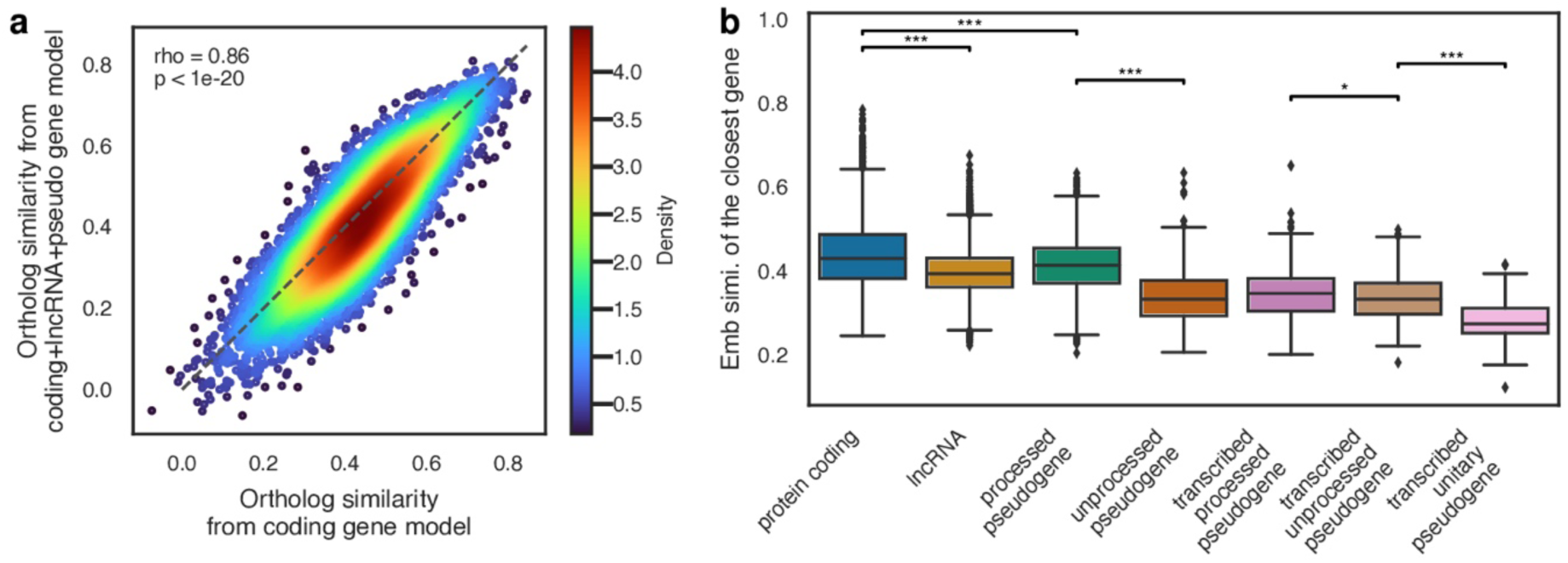
Embedding similarities from the coding genes, lncRNAs and pseudogene model. **a,** Spearman correlation of one-to-one orthologous gene embedding similarities from the protein-coding gene-only model and the model including coding genes, lncRNAs and pseudogenes. The dashed line represents the line of equality (y=x). **b,** Distribution of embedding similarity for the closest gene across different gene groups. Gene embeddings from the model of coding genes, lncRNAs, and pseudogenes were used. In each gene group, for each human gene, the plot displays the similarity with its closest mouse gene on the vertical axis. Note that genes used for supervision in the supervised embedding alignment were excluded from the protein-coding gene group in the plot. Boxplot details: center line indicates the median; box limits represent the upper and lower quartiles; whiskers extend to 1.5× the interquartile range; points beyond the whiskers are outliers. In boxplots, two-sided Wilcoxon rank-sum test was used for comparisons. *** p < 0.00001, ** p < 0.001, * p < 0.05, ‘ns’ denotes not significant (p ≥ 0.05).

Next, we classified the genes into seven groups: protein-coding genes, lncRNAs, and five pseudogene groups (Fig. 4b). For each group, we identified the most similar mouse gene for each human gene based on gene embedding. To avoid bias, orthologs used for supervision in supervised alignment were excluded when assessing similarity. The analysis revealed that protein-coding genes had the highest similarity between the two species (Fig. 4b), suggesting a conservation of their characteristics at the RNA level. Processed pseudogenes and lncRNAs followed, and the other pseudogene groups had the lowest similarity, indicating more expression-level variability between species. These RNA similarity insights could be valuable for understanding gene conservation at the RNA level, and potentially provide new information about gene functions and attributes. Furthermore, they could assist in identifying appropriate mouse representative genes for human lncRNAs and pseudogenes in biomedical research.

## Discussion

In this study, we developed a novel deep learning approach to characterize and compare human and mouse genes at the RNA level, using a large bulk RNA-seq dataset. Our thorough optimization and evaluation of gene embedding alignment significantly improves the methodology and offers valuable lessons for future studies. Our RNA-level representations offer insights beyond DNA sequence similarity, aiding in selecting appropriate animal models for gene studies and improving our understanding of genes in humans and mice.

Aligning gene embeddings from different species into the same space opens up the possibility of using advanced deep learning technology for inter-species comparisons. This approach could potentially be extended to aligning different biological conditions and even different omics data. Our experiments demonstrate that mixed training, shared embeddings, and supervised alignment techniques can improve alignment quality. Importantly, our study found that alignment results were robust to the alignment techniques used, making this a powerful approach even when some aligning techniques may not be applicable in certain future scenarios.

The inter-species DNA and RNA similarities of genes do not fully overlap. Some homologous gene pairs exhibited low DNA similarity but high RNA similarity, suggesting that despite differences in DNA sequences, they still have similar expression relationships with other genes. On the other hand, some gene pairs had similar DNA sequences but low RNA similarity, suggesting that their functions may have diverged. Therefore, we suggest exercising caution when selecting animal models or interpreting results from mouse models for studying genes with low RNA similarity, particularly those with low similarity in both DNA and RNA.

It’s important to note that our RNA similarities reflect the average across tissues and biological conditions. In certain applications, it may be beneficial to use the RNA similarity for a specific tissue or a specific biological condition. While not explicitly tested, it is theoretically feasible to obtain tissue- or condition-specific similarities by adding corresponding expression embeddings to the gene embeddings, as this is the operation adopted by the Transformer model during input, and our GeneRAIN study provided preliminary results supporting this method^12^. Although our similarities are the averaged similarities, they still provide valuable information about gene function. Furthermore, our averaged similarities can be helpful for studying the evolution of the two species and potentially enhance our understanding of their gene biology.

## Methods

### Data preprocessing

Gene-level human and mouse bulk RNA-seq data were retrieved from the ARCHS4 database (version 2.2)^38^. Data collection and processing adhered to ethical guidelines, under the supervision and approval of the University of New South Wales (UNSW) Human Research Ethics Committee. The data processing followed the methodology described in our previous GeneRAIN study^12^. Briefly, sample selection was based on stringent criteria: total gene expression read counts exceeding one million, a minimum of 2,000 genes exhibiting non-zero expression, and a single-cell probability below 0.5 (as estimated by the database) to eliminate potential single-cell samples. These criteria ensured the inclusion of high-quality samples.

The samples underwent library size normalization to have a uniform total read count of 10 million per sample. Subsequently, read counts were log-transformed (base 10), and only genes with a mean normalized expression above 0.1 were retained. This selection process resulted in a final dataset comprising 18,762 human protein-coding genes, 13,006 human lncRNA genes, 4,956 human pseudogenes, 19,072 mouse protein-coding genes, 7,178 mouse lncRNA genes, and 3,599 mouse pseudogenes.

Expression data discretization followed the procedure outlined in our previous GeneRAIN study. Gene expressions across all samples were categorized into 2,000 bins according to their expression ranks, using the Binning-By-Gene normalization method. This approach ensured that each bin, except the very first, contained equal number of samples. Before being fed into the Transformer GPT^37^ or BERT^36^ models, gene expressions in each sample were ordered from highest to lowest based on their bin assignments. The models received only the top 2048 genes, with attention paying to any genes in the inputs, including those with zero expression. In the BERT model specifically, 15% of genes exhibiting non-zero expression were randomly selected to have their identities (gene symbols) masked. The model was trained to reconstruct the identities of those genes.

### Gene annotations

DNA sequence data was downloaded from Table Browser of UCSC Genome Browser^57^. For genic region annotation and gene transcript annotation, the GENCODE database was used^46^. Specifically, for humans, the GRCh38 assembly and GENCODE V44 were used. For mice, GRCm39 and GENCODE VM33 were used. Gene type annotations, such as annotations of being protein coding genes, lncRNA genes, etc., were primarily extracted from the GENCODE database. For genes whose information was not available in GENCODE database, gene type annotations from the metadata in the ARCHS4 data file was used. The information for mapping transcript IDs to gene names was obtained from Ensembl database Release 111 (https://ftp.ensembl.org/pub/)^58^.

Information regarding human-mouse homologs and orthologs was obtained from the file ‘HOM_MouseHumanSequence.rpt’ in the MGI database^40^, downloaded in January 2024. One-to-one homologs identified in this file were used as one-to-one orthologs.

### Model hyperparameters

The configuration and parameters of the model were consistent with those detailed in our previous GeneRAIN GPT study. Briefly, each gene and its expression value were represented in a 200-dimensional space. In both the BERT and GPT models, gene embedding and expression embedding were added together. The architecture of the model consisted of six layers, with each layer containing four attention heads and each head had 32 dimensions. The AdamW optimizer was utilized for model optimization^59,60^, starting with a base learning rate of 0.00001 and allowing it to peak at 0.0001. To optimize memory utilization and manage batch size effectively, gradient accumulation steps were set at 5. The batch size was 12 for coding genes only model, and batch size was set to 8 for models with coding gene, lncRNAs and pseudogenes due to memory constraint. The dataset was divided into training (90%) and validation (10%) sets.

### Gene embedding alignment

In the mixed training approach, human samples were randomly mixed with mouse samples to form the training dataset (Fig. 1a). Each mouse gene symbol was prefixed with “m_” to distinguish it from human gene symbols. While human genes had gene embeddings different from those of mouse genes, both shared the same expression embeddings. Each training instance was either a single human sample or a single mouse sample.

For the shared embedding training, three separate models were trained. Within each model, 5,000 one-to-one orthologous gene pairs were randomly selected, and each selected orthologous gene pair, consisting of a human gene and a mouse gene, was assigned the same gene embedding. Importantly, the selection of 5,000 orthologous gene pairs for each model was non-overlapping, ensuring that none of those gene pairs were selected in more than one model. After training, we only used information from the genes that did not have shared embeddings. The cosine similarity between these genes was computed using their respective embeddings from each model. We then averaged the similarities from the three models, and used them as the final similarity values from this embedding alignment approach.

For unsupervised alignment, the program ‘unsupervised.py’ implemented by the MUSE python package was used (https://github.com/facebookresearch/MUSE)^42^. The training was conducted for 5 epochs. The function for selecting embeddings of the k most frequent words for discrimination was disabled, and other parameters were set to their default values.

In the case of supervised alignment, the program ‘supervised.py’ implemented by the MUSE python package was employed (https://github.com/facebookresearch/MUSE). A random selection of 95% (n=16134) of one-to-one orthologous gene pairs constituted the training set, while the remaining 5% (n=849) served as the evaluation set. Five refinement iterations were performed. All other parameters were maintained at their default settings.

For the shared embedding plus supervised alignment approach, three shared embedding models were first trained as previously described. Subsequently, supervised alignment was applied to the genes without shared embeddings in each model. We calculated the average gene-pair embedding similarities across the three models, and used them as the final similarities from this approach.

### Evaluation of embedding alignment

In the pseudo non-human ‘positive control’ experiment, we created a pseudo non-human species. The 410K human RNA-seq samples were divided into two halves, each comprising 205K samples. The first 205K samples were labeled as real human samples, and the remaining 205K samples were assigned to the pseudo non-human species. New gene symbols were ‘invented’ for each gene in the pseudo non-human species, and the mapping relationship with the actual human gene symbols was documented. That relationship was later used for embedding alignment supervision and evaluation of the alignment performance.

Different embeddings were assigned to the genes in the real human species and the genes in the pseudo non-human species. This resulted in embeddings for 18,757 genes from the real human species and another set of embeddings for 18,757 ‘invented’ genes from the pseudo non-human species. The real human and pseudo non-human species underwent mixed training. Post mixed training, the documented mapping relationship between the actual human gene symbols and the ‘invented’ genes from the pseudo non-human species was used for supervision to align the embeddings from these two ‘species’. Subsequently, we assessed the extent to which the mapping relationships were reconstructed by the alignment.

In the shuffled embeddings ‘negative control’ experiment, real human samples and real mouse samples first underwent mixed training as normal. Then prior to supervised alignment, the order of embeddings within each species was randomly shuffled, thereby disrupting the gene-embedding correspondence. Supervised alignment, utilizing the one-to-one orthologous gene pairs for supervision, was conducted on these shuffled embeddings. Subsequently, we assessed the extent to which the one-to-one orthologous relationships were reconstructed by the alignment.

For evaluating embedding alignment methodologies and experiments, the ranks of ortholog embedding similarity were examined. The process was performed as follows: for each human gene with a corresponding one-to-one mouse ortholog, its cosine similarity with every mouse gene was calculated using gene embeddings. Subsequently, all mouse genes were ranked based on these similarities, from highest to lowest. The rank of the corresponding mouse ortholog for the human gene was recorded. This procedure was repeated for every human gene (if a one-to-one mouse ortholog is available). The ranks for all the human genes were then used for assessing the closeness of embeddings between orthologous genes, and for evaluating the performance of the models or alignment methods.

Additionally, the ranking process could be reversed, starting with each mouse gene and examining the ranks of its human orthologs. Evaluations indicated that rankings in both directions are highly correlated, as detailed in Extended Data Fig. 5e, f.

### DNA alignment

For the DNA alignment, the Biopython package was employed, specifically using PairwiseAligner function from Bio.Align module^61^. Global alignment was performed at the transcript level for each gene. Transcripts with a length of less than 100 base pairs (bp) were excluded from the analysis. For our pairwise alignments, following scoring system was used, the match score was set to 2, penalizing mismatches and gap openings with scores of −1 and −2, respectively, and a gap extension penalty of −1. The calculated scores for a pairwise alignment were normalized by the average length of the two sequences being compared, to facilitate the comparison of alignment scores across different gene pairs.

For aligning sequences from two genes, such as Gene A from humans and Gene B from mice, following approached was used. For each transcript in Gene A, we identified the best possible matched transcripts by finding the transcript of Gene B that result in highest alignment. Subsequently, the maximum of scores from all transcripts in Gene A were used as the DNA alignment score between Gene A and Gene B. While the mean of scores from all transcripts in Gene A were used as the average transcript alignment score. This approach provided a measure of similarity, taking into account the splicing diversity of genes.

### Phenotype association analysis

For our human and mouse gene phenotype association analysis, we used the ‘HMD_HumanPhenotype.rpt’ file from the the ‘Mouse/Human Orthology with Phenotype Annotations’ section in the Mouse Genome Informatics (MGI) database (https://www.informatics.jax.org/downloads/reports/index.html)^40^. From this file, we extracted the number of high-level mammalian phenotypes associated with each gene pair. The extracted numbers were then used for comparative analysis and visualization.

For human disease mouse model association analysis, we obtained the annotation data from the ‘MGI_DiseaseGeneModel.rpt’ file, which located in the ‘Mouse models of Human Disease by Human gene’ section in the MGI database. This file provided information on mouse models corresponding to human diseases. For each gene pair, we counted the number of their associated ‘DO term names’, for comparative analysis and visualization.

For Alliance disease association analysis, we obtained the annotation dataset from the Alliance of Genome Resources website, Version: 6.0.0 (https://www.alliancegenome.org/downloads)^45^. Specifically, we used two compressed tab-separated values (tsv) files: ‘DISEASE-ALLIANCE_HUMAN.tsv.gz’ for human genes and ‘DISEASE-ALLIANCE_MGI.tsv.gz’ for mouse genes. This dataset provided comprehensive information on gene-disease associations for both human and mouse genes. We extracted gene symbols and associated disease names from these files using columns ‘DBObjectSymbol’ and ‘DOtermName’, respectively. Subsequently, as association type and evidence type information is available in this dataset, we performed filtering to include only well-supported association records.

For human genes, we included association records with ‘AssociationType’ being ‘is_implicated_in’ or ‘is_marker_for’, and excluded those being ‘implicated_via_orthology’, ‘biomarker_via_orthology’, or ‘is_not_implicated_in’. Additionally, we applied filtering to the association evidence. We selected only records where the ‘EvidenceCodeName’ included ‘inference by association of genotype from phenotype used in manual assertion’, ‘direct assay evidence used in manual assertion’, and ‘mutant phenotype evidence used in manual assertion’, excluding those supported only by automatic assertions.

Similarly, for mouse genes, we selected records with ‘DBobjectType’ being ‘gene’ (excluded ‘allele’ and ‘affected_genomic_model’) and ‘AssociationType’ being ‘is_implicated_in’ or ‘is_marker_for’. We also applied filtering to the ‘EvidenceCodeName’, retaining only those being ‘author statement supported by traceable reference’. These filters provided us a dataset of high-confidence association records, enabling us to perform association and comparative analyses on homologs with different DNA and RNA similarities.

### Comparison with LECIF scores

To obtain LECIF scores^33^, we downloaded the bigwig format^62^ file ‘hg38.LECIFv1.1.bw’ from https://github.com/ernstlab/LECIF and converted it to wig format using the UCSC bigWigToWig tool^57^. Since LECIF scores are assigned to 50-bp bins, we calculated gene-level LECIF scores to facilitate phenotype and disease association comparisons. This was done by averaging LECIF scores for all positions within each gene’s genic region. We also calculated a separate set of scores using only positions within coding regions. Genomic coordinates for both genic and coding regions were obtained from GENCODE V44 annotation^46^. We then compared the ability of both versions of calculated LECIF scores and our embedding similarities in predicting whether homologous gene pairs shared phenotype(s) or disease association(s).

### Data availability

The supervised aligned gene embeddings derived from the protein-coding genes-only GPT model, as well as from the model of coding genes, lncRNAs + pseudogenes, are available at https://zenodo.org/records/10866876. These embeddings allow for straightforward calculation of pairwise similarity. The 37,926 by 37,926 pairwise similarity matrix, produced by averaging the similarities from the supervised alignment approach and those from shared embeddings + supervised alignment approach, is also available at the same link. Provided files are made available under the Creative Commons Attribution Non Commercial 4.0 International License. All data is free to use for non-commercial purposes.

### Code availability

The source code used for training the models, aligning cross-species gene embeddings, and performing all the analyses to generate the results in the study, is available in GitHub repository https://github.com/suzheng/GeneRAIN_HM. Provided code is made available under the Creative Commons Attribution Non Commercial 4.0 International License. All code is free to use for non-commercial purposes.

## Acknowledgements

We would like to thank the Research Technology Services (ResTech) at the University of New South Wales for their help with computational resources, with a special thank you to Martin Thompson for his support and assistance. We also extend our thanks to the National Computational Infrastructure (NCI) for their support. This research was undertaken with the assistance of resources and services from the National Computational Infrastructure (NCI Australia), which is supported by the Australian Government through the National Collaborative Research Infrastructure Strategy (NCRIS). Z.S. acknowledges funding support from The University of New South Wales (University Postgraduate Award scholarship).

## Author contributions

Z.S. and M.F conceptualized and designed the research, Z.S. conducted the data analysis. E.O., M.D. and F.V. provided project supervision and strategic guidance on research methodologies and analysis. Z.S. and M.F. generated visualizations, and led the writing of the manuscript. All authors contributed to the intellectual content, provided critical feedback, participated in revising the manuscript, and approved the final version for publication.

## Extended Data

**Extended Data Fig. 1.**
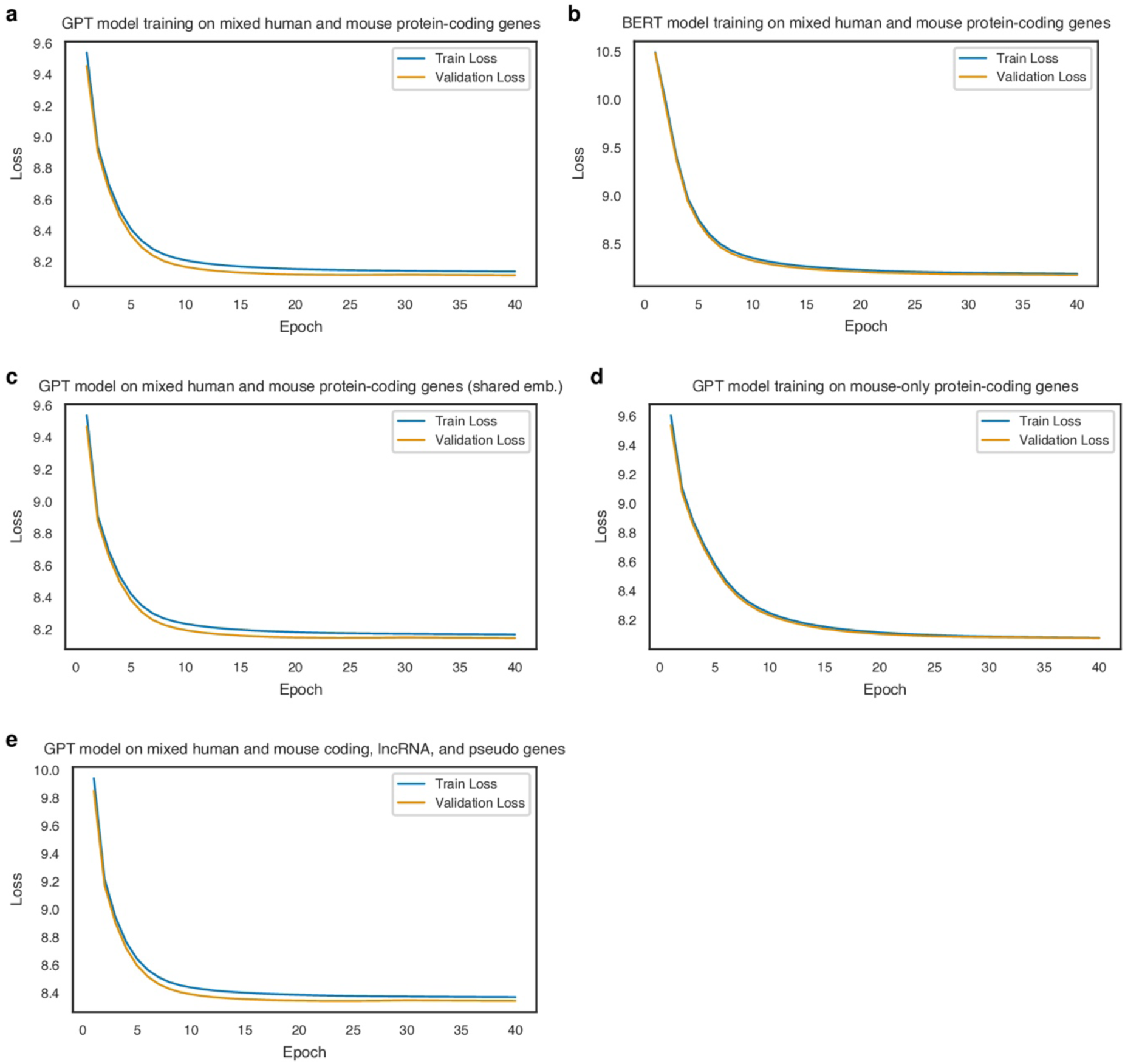
Training and validation loss of across epochs. This figure illustrates the training and validation loss over epochs for various models.

**Extended Data Fig. 2.**
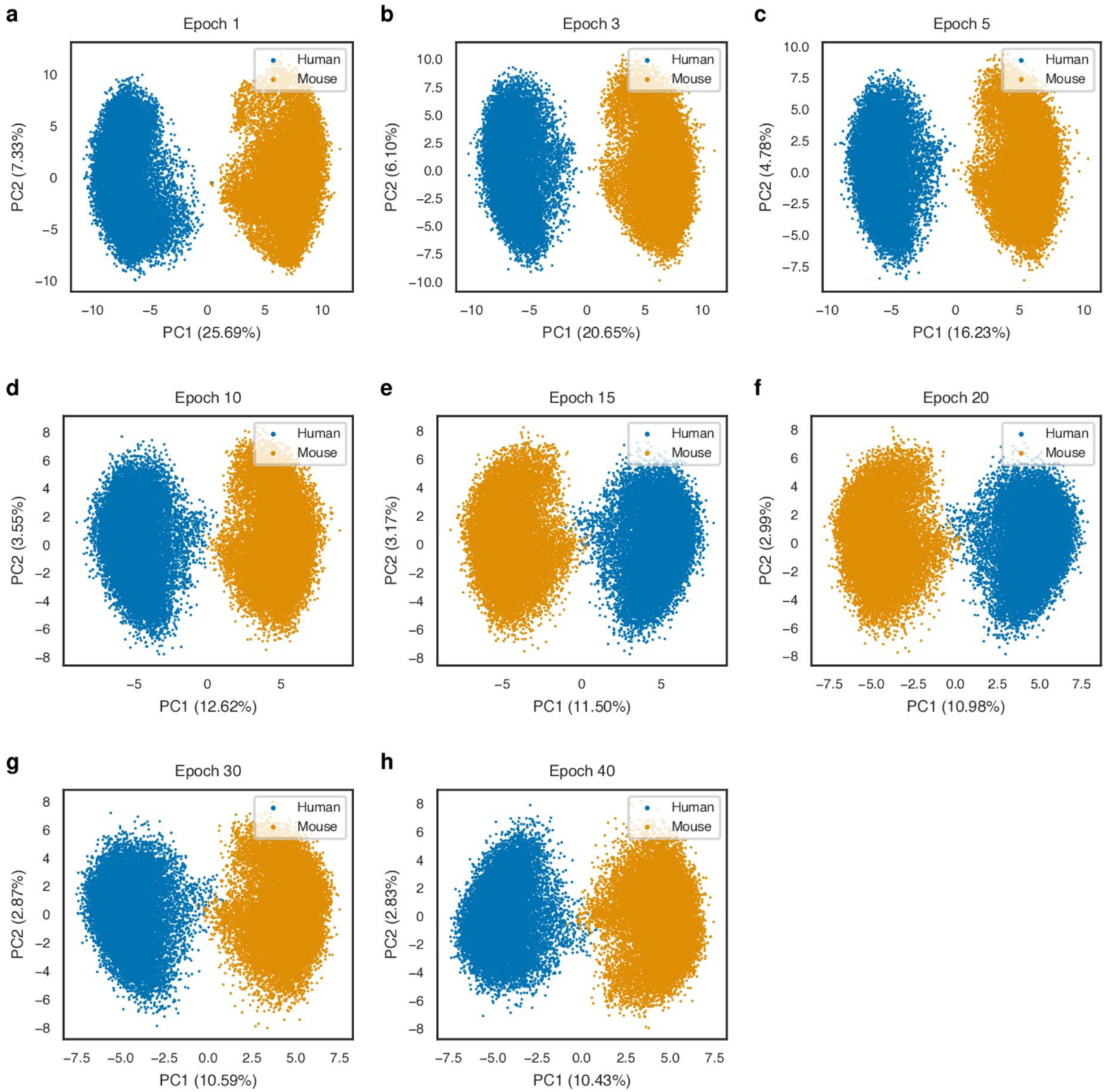
PCA plots of gene embedding distributions over training epochs. The plots illustrate the distribution of gene embeddings derived from the mixed training GPT model at various training epochs. Each dot in the plots represents a gene. Please note the percentage of explained variance on the x-axis.

**Extended Data Fig. 3.**
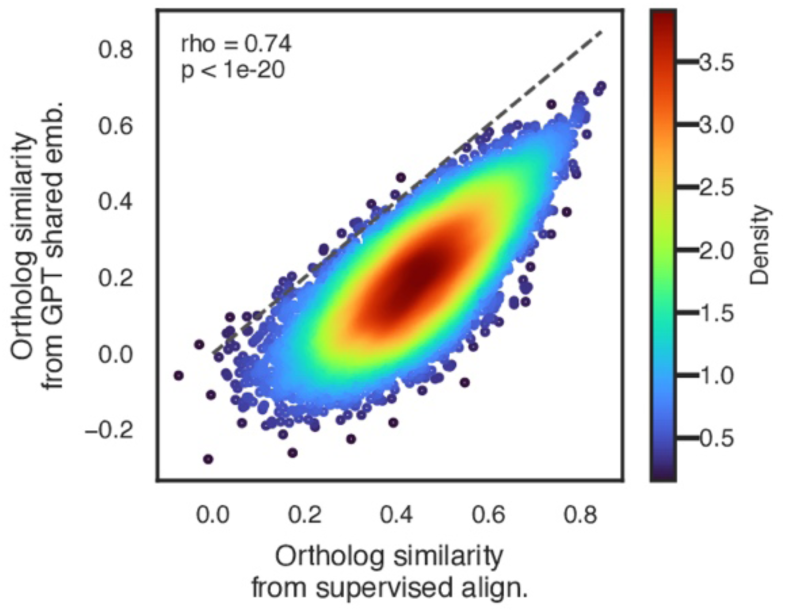
Embedding similarities from the shared embedding and supervised alignment approaches. This scatter plot visualizes the Spearman correlation of orthologous gene embeddings similarities from these two embedding alignment approaches. The dashed line represents the line of equality (y=x).

**Extended Data Fig. 4.**
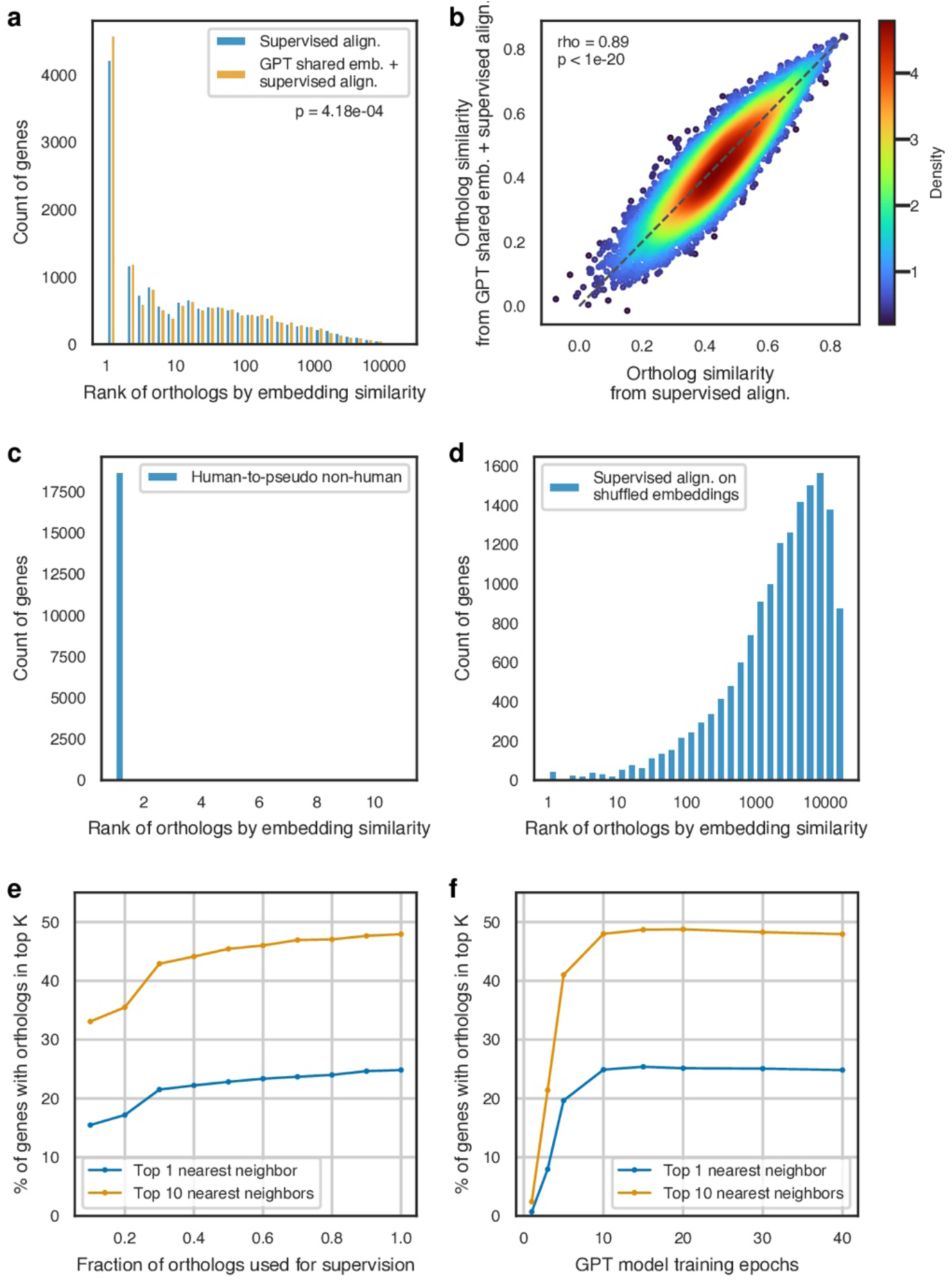
Evaluation of embedding alignment strategies. **a,** Distribution of embedding similarity ranks for one-to-one orthologs from supervised alignment only and shared embedding plus supervised alignment approaches, with statistical significance evaluated by a two-sided Kolmogorov-Smirnov test. **b,** Spearman correlation of one-to-one orthologous gene embedding similarities from these two alignment approaches. The dashed line represents the line of equality (y=x). **c,** Distribution of embedding similarity ranks for one-to-one orthologs from the pseudo non-human species in the ‘positive control’ experiment. **d,** Distribution of embedding similarity ranks for orthologs from the shuffled embedding ‘negative control’ experiment. **e,** Efficiency of supervised alignment with varying fractions of orthologous gene pairs used for supervision, illustrated by the percentage of human genes with their mouse orthologs ranking as the closest gene (blue line), and within the top 10 genes based on embedding similarity. **f,** Efficiency of supervised alignment across different numbers of training epochs.

**Extended Data Fig. 5.**
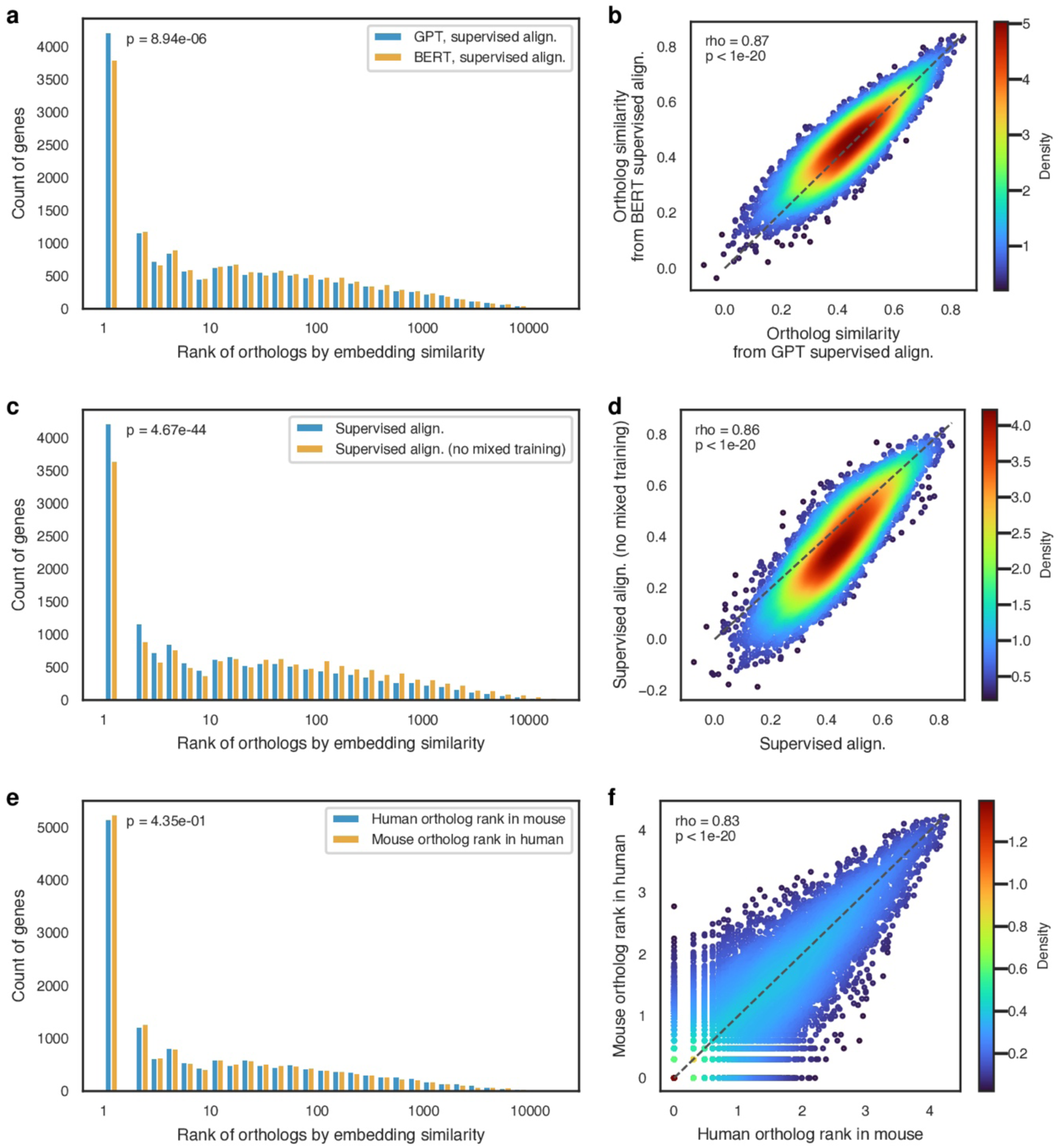
Evaluation of embedding from different models. **a,** Distribution of embedding similarity ranks for one-to-one orthologs from GPT and BERT models, with mixed training and supervised alignment were used for both models. **b,** Spearman correlation of one-to-one orthologous gene embedding similarities from the GPT and BERT models. **c,** Distribution of embedding similarity ranks for one-to-one orthologs from models with and without mixed training. GPT model and supervised alignment were applied in both scenarios. **d,** Spearman correlation of one-to-one orthologous gene embedding similarities from the models with and without mixed training. **e,** Comparative analysis of human ortholog ranks in two directions. The rank of each human gene’s ortholog within all mouse genes was calculated using embeddings from the shared embedding plus supervised alignment approach. Similarly, the rank of each mouse gene’s ortholog within all human genes was also determined. This histogram visualizes the distribution of these two sets of ranks. **f,** A scatter plot showing the correlation between these two sets of ranks, assessed using Spearman’s rho and p-value. In all barplots, statistical significance was evaluated by a two-sided Kolmogorov-Smirnov test. In all scatter plot, each dot symbolizing a orthologous gene pair. The dashed line represents the line of equality (y=x).

**Extended Data Fig. 6.**
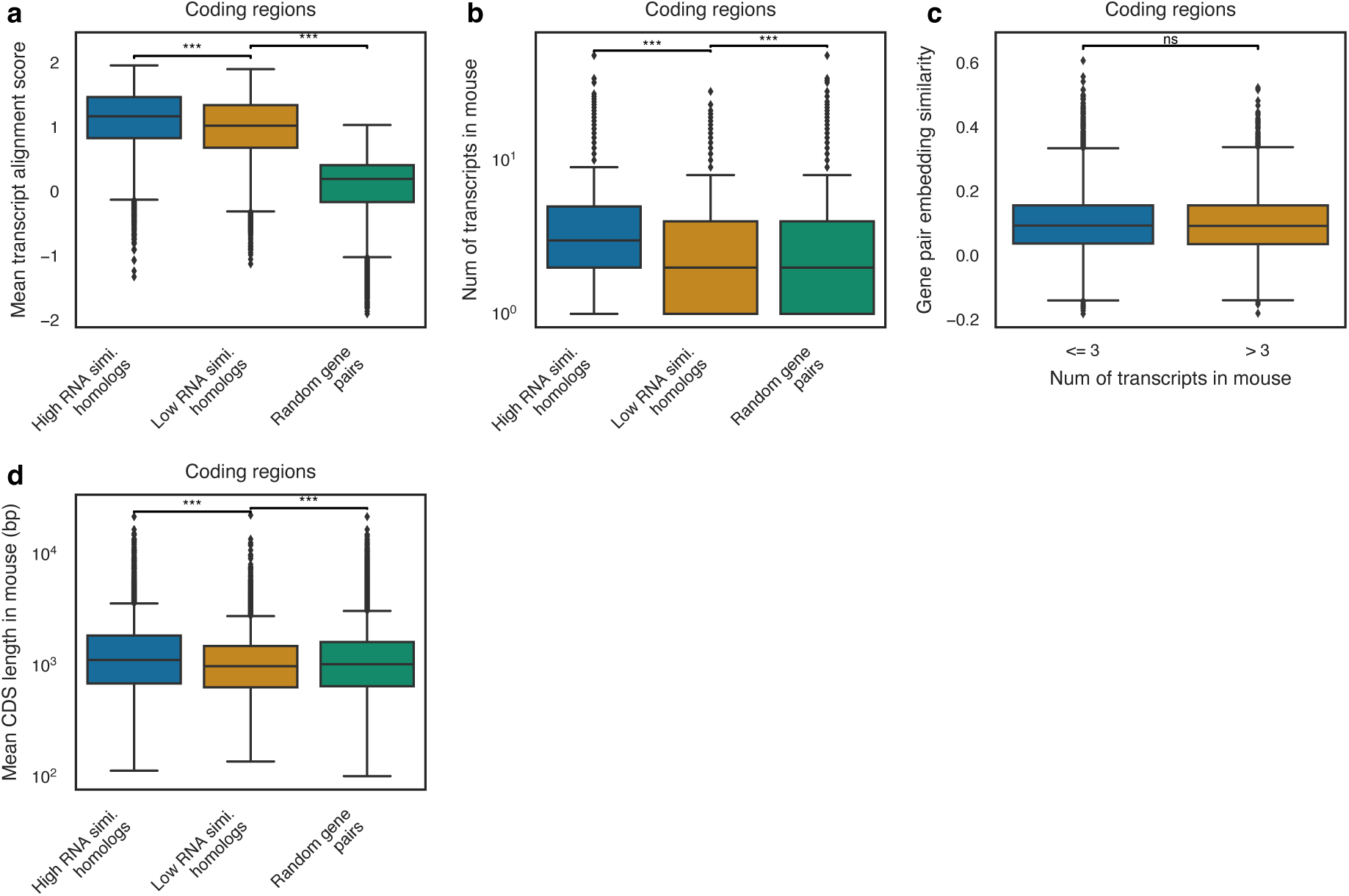
DNA characteristics of homologs grouped by embedding similarity. **a-c,** Boxplots depicting mean transcript alignment scores (Methods) for **(a)** coding sequences, **(b)** 1kb upstream regions of transcription start sites, and **(c)** 1kb downstream regions of transcription termination sites in homologs with high embedding similarity (n = 7,249) and low embedding similarity (n = 7,103). Mean transcript alignment scores are calculated as the average pairwise alignment similarity across the human transcripts for each gene pair (Methods), offering a complementary perspective on DNA sequence conservation. Random gene pairs are included for comparison. **d,** Average coding sequence length in the human genes from three gene pair groups. In all boxplots: center line, median; box limits, upper and lower quartiles; whiskers, 1.5× interquartile range; outliers, points. All comparisons were evaluated using a two-sided Wilcoxon rank-sum test. *** *p* < 0.00001, ** *p* < 0.001, * *p* < 0.05, ‘ns’ not significant (*p* >= 0.05).

**Extended Data Fig. 7.**
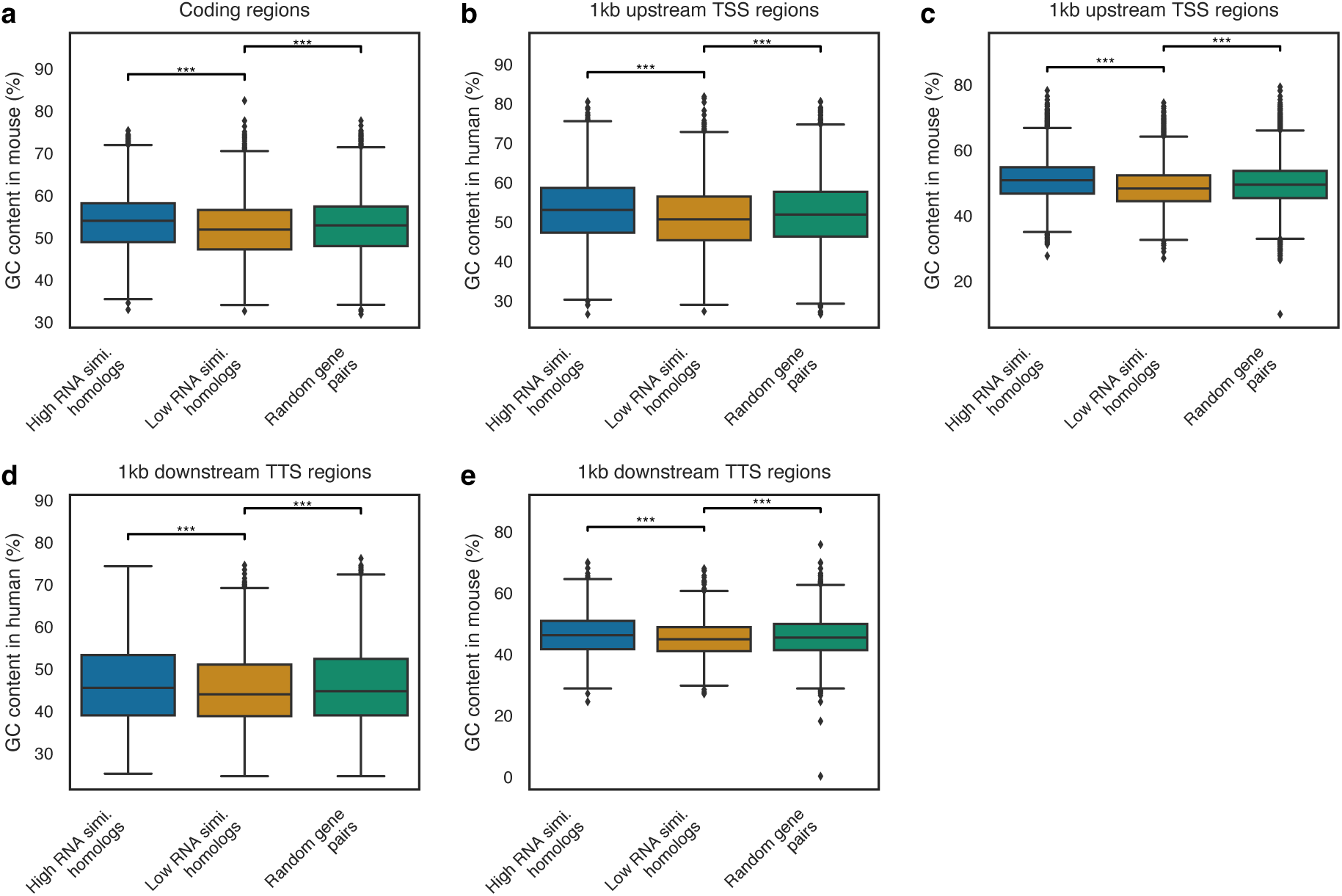
GC content of homologs grouped by embedding similarity. **a,** Boxplots visualize the GC content of the coding sequences of mouse genes from homologs with high embedding similarity (n = 7,249) and low embedding similarity (n = 7,103). Random gene pairs are included for comparative analysis. **b-e,** Analysis of GC content in regulatory regions. With **b** for the 1kb upstream of transcription start sites in human genes; **c** for the 1kb upstream of transcription start sites in mouse genes; **d** for the 1kb downstream of transcription termination sites in human genes; and **e** for the 1kb downstream of transcription termination sites in mouse genes. Each analysis compares the two homolog groups and a random gene pair group. In all boxplots: center line, median; box limits, upper and lower quartiles; whiskers, 1.5× interquartile range; outliers, points. All comparisons were evaluated using a two-sided Wilcoxon rank-sum test. *** *p* < 0.00001, ** *p* < 0.001, * *p* < 0.05, ‘ns’ not significant (*p* >= 0.05).

**Extended Data Fig. 8.**
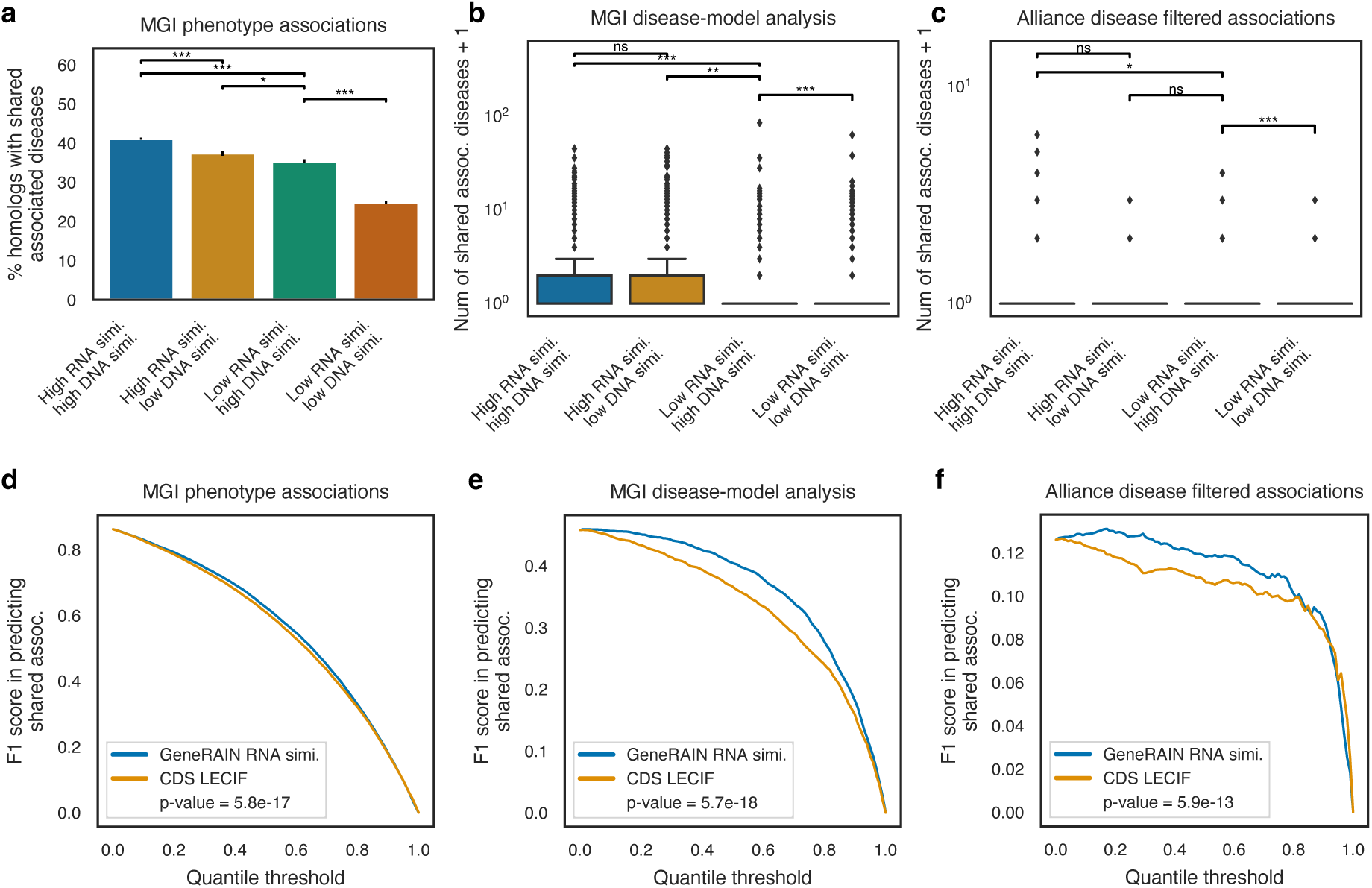
Embedding similarity and phenotype association analysis. **a** Barplot displaying the percentages of homologs with shared associated phenotype associations, based on ortholog phenotype association annotations from MGI database. Homologs were categorized by DNA similarity and embedding similarity learned from RNA data. Error bars represent the standard errors. **b,** Boxplot for number of shared associated diseases across homolog groups, according to the human disease mouse model annotations from MGI. **c,** Barplot showing number of shared associated diseases across homolog groups, according to filtered disease association annotations from the Alliance of Genome Resources database (Methods). In all boxplots: center line, median; box limits, upper and lower quartiles; whiskers, 1.5× interquartile range; outliers, points. In barplots, two-sample z-test for proportions was used for comparisons, while in boxplots, two-sided Wilcoxon rank-sum test was used. *** *p* < 0.00001, ** *p* < 0.001, * *p* < 0.05, ‘ns’ not significant (*p* >= 0.05). **d-f**, F1 Score vs. threshold curves comparing the performance of our RNA similarities and LECIF scores in predicting whether homologous gene pairs have shared association(s), using annotations from the database as indicated in the title. For LECIF, in contrast to Fig. 3, the average score within the coding region rather than genic region was used for each gene. The curves visualize how the F1 score (harmonic mean of precision and recall) varies across different classification thresholds. The p-values, obtained from the Wilcoxon signed-rank test, compare the F1 scores of the two metrics across 100 quantile thresholds.

