## Supplementary Information for "Genes in Humans and Mice: Insights from Deep learning of 777K Bulk Transcriptomes"

**Optimization and evaluation of gene embedding alignment**

We investigated whether combining shared embeddings with supervised alignment methods could further enhance the performance. This experiment involved applying supervised alignment to genes with non-shared embeddings from the shared embedding models. The comparative analysis indicated a modest improvement (4,573 vs 4,216 orthologs being the closest), particularly when considering the 10 closest embeddings (8,255 vs 8,140) (Extended Data Fig. 4a). The embedding similarities from two approaches demonstrated a strong correlation (Spearman correlation rho = 0.89, p < 1e-20, Extended Data Fig. 4b).

In our previous analysis, fewer than 5,000 of 16,983 mouse genes were the nearest embedding neighbor of their human orthologs. To investigate if this was due to limitations of the methodology, we conducted a ‘positive control’ experiment. We invented a pseudo non-human species, along with its own gene symbols and gene embeddings. We then divided 410K human RNA-seq samples into two halves, one for the real human and another for the pseudo species. Then the two ‘species’ underwent normal mixed training and supervised alignment processes (Methods). We found that 18,695 of 18,757 (99.7%) genes had their inter-‘species’ relationships reconstructed by the model (Extended Data Fig. 4c). This indicates the high effectiveness of our approach.

We then assessed the false positive rate of our supervised alignment approach, addressing the concern that it might arbitrarily align genes in the supervision set, regardless of their embeddings. For that assessment, we conducted a 'negative control' experiment where we randomized the embedding order after mixed training, disrupting the gene-embedding correspondence. This randomized setup was then subjected to supervised alignment. In this experiment, only 45 of 16983 (0.3%) of orthologs had closest embedding (Extended Data Fig. 4d), indicating minimal spurious appearance of relationship used in supervision.

We then investigated how alignment outcomes can be affected by training parameters, including the number of orthologous gene pairs used for supervision and training epochs. Analysis showed that the result started to plateau with just 50% of orthologous gene pairs used for supervision (Extended Data Fig. 4e) or after 10 epochs of training (Extended Data Fig. 4f). We then investigated the influence of model architectures, strong correlation between results from GPT and Bidirectional Encoder Representations from Transformers (BERT) model^34^ was observed (Spearman correlation rho = 0.87, p<1e-20), with slightly superior performance in GPT (Extended Data Fig. 5a, b). Finally, we assessed supervised alignment with and without mixed training, we found mixed training produced better alignment (Extended Data Fig. 5c, d), underscoring its value in aligning gene embeddings.

Beside gene similarity values, we also used the ranks of embedding closeness to quantify inter-species gene relationships. Further, we compared the ranks in opposite directions (mouse-to-human versus human-to-mouse), we found that they were highly correlated (Extended Data Fig. 5e, f).
